# The spatial and cell-type distribution of SARS-CoV-2 receptor ACE2 in human and mouse brain

**DOI:** 10.1101/2020.04.07.030650

**Authors:** Rongrong Chen, Keer Wang, Jie Yu, Derek Howard, Leon French, Zhong Chen, Chengping Wen, Zhenghao Xu

**Author notes:** Corresponding Author: Chengping Wen, PhD, MD. Binwen Road 548, Hangzhou, Zhejiang, China, Zhenghao Xu, PhD. Binwen Road 548, Hangzhou, Zhejiang, China. The first three authors contribute equally.

## Abstract

By engaging angiotensin-converting enzyme 2 (ACE2 or Ace2), the novel pathogenic SARS-coronavirus 2 (SARS-CoV-2) may invade host cells in many organs, including the brain. However, the distribution of ACE2 in the brain is still obscure. Here we investigated the ACE2 expression in the brain by analyzing data from publicly available brain transcriptome databases. According to our spatial distribution analysis, ACE2 was relatively highly expressed in some brain locations, such as the choroid plexus and paraventricular nuclei of the thalamus. According to cell-type distribution analysis, nuclear expression of ACE2 was found in many neurons (both excitatory and inhibitory neurons) and some non-neuron cells (mainly astrocytes, oligodendrocytes, and endothelial cells) in human middle temporal gyrus and posterior cingulate cortex. A few ACE2-expressing nuclei were found in a hippocampal dataset, and none were detected in the prefrontal cortex. Except for the additional high expression of Ace2 in the olfactory bulb areas for spatial distribution as well as in the pericytes and endothelial cells for cell-type distribution, the distribution of Ace2 in mouse brain was similar to that in the human brain. Thus, our results reveal an outline of ACE2/Ace2 distribution in the human and mouse brain, which indicates the brain infection of SARS-CoV-2 may be capable of inducing central nervous system symptoms in coronavirus disease 2019 (COVID-19) patients. Potential species differences should be considered when using mouse models to study the neurological effects of SARS-CoV-2 infection.

## Introduction

Since December 2019, much attention has focused on the novel SARS-coronavirus 2 (SARS-CoV-2) and related coronavirus disease 2019 (COVID-19), which is rapidly spreading around the world and resulted in a global health emergency ^[1]^. In addition to atypical pneumonia, the central nervous system (CNS) symptoms of COVID-19 patients have been observed in the clinic ^[2]^. According to a recent retrospective case series study, 53 out of 214 (24.8%) COVID-19 patients had CNS symptoms, including dizziness, headache, impaired consciousness, acute cerebrovascular disease, ataxia, and epilepsy ^[3]^. More importantly, it has been found that SARS-coronavirus (SARS-CoV), a previous similar coronavirus to SARS-CoV-2, spread into the brain after it was cleared from the lung in mice, which could be more concealment than that in the lung ^[4]^. Recently, Puelles et al. found that SARS-CoV-2 has an organotropism beyond the respiratory tract, including the kidneys, liver, heart, and brain^[5]^. Thus, it is necessary and urgent to pay attention to the CNS infection of SARS-CoV-2.

Angiotensin-converting enzyme 2 (ACE2 or Ace2) has been identified as a key entry receptor for novel pathogenic SARS-CoV-2 as well as previous SARS-coronavirus (SARS-CoV) ^[6]^. By binding of the spike protein of the virus to ACE2, SARS-CoV-2 and SARS-CoV could invade host cells in human organs ^[7, 8]^. However, the distribution of ACE2 in the brain is still obscure and even inconsistent. Hamming et al. found ACE2 may be express only in endothelium and vascular smooth muscle cells in the human brain tissue in 2004 ^[9]^. Since then, few studies focused on the distribution of ACE2 in the human brain. On the other hand, a previous study has reported that Ace2 could express in mouse neuron cells, which may contribute to the development of hypertension ^[10]^; however, in another neurocytometry study, Ace2 is a potential marker for non-neurons in zinc-fixed mouse brain cortical section ^[11]^. Thus, further clarifying brain tissue distribution of ACE2 may help to bring light to the CNS infection of the novel SARS-CoV-2 and previous SARS-CoV.

Here, we investigated the distribution of ACE2 in the brain by analyzing publicly available brain transcriptome databases. We revealed an uneven spatial and cell-type distribution of ACE2 in the human and mouse brain.

## Methods

### Brain transcriptome databases

Seven publicly available brain transcriptome databases that can be accessed without specialized computational expertise were used. All databases were listed in Table 1. Except for the Single Cell Portal database (https://singlecell.broadinstitute.org), these databases have been also introduced in a recent review study ^[12]^. All databases and datasets were appropriately used and cited according to their citation policy, license, or terms of use.

**Table 1.**
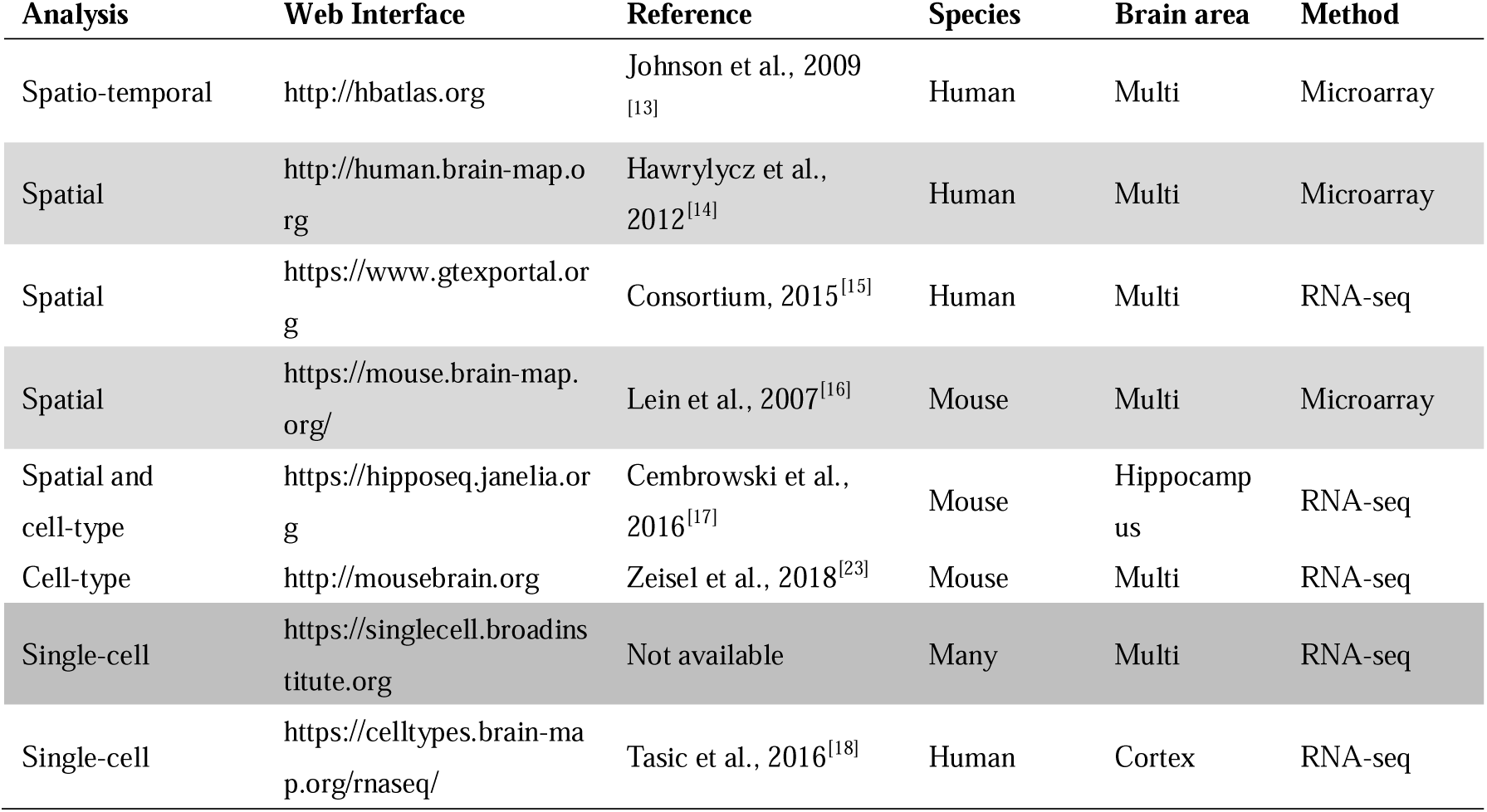
The database used for the current study.

### Analysis of the spatial distribution of ACE2 in the human brain

Three databases were used, including Allen human brain atlas database (http://human.brain-map.org), Human Brain Transcriptome database (https://hbatlas.org), and GTExportal database (https://www.gtexportal.org), to analyze the spatial distribution of ACE2 in the human brain.

### Cell-type distribution of ACE2 in the human brain

Two single-cell sequencing databases, including Single Cell Portal database (https://singlecell.broadinstitute.org, Single-cell sequencing) and Allen Cell Types Database (http://celltypes.brain-map.org, Single-cell sequencing), were used. The summary of the included three datasets for the human brain were shown in Table 2.

**Table 2.**
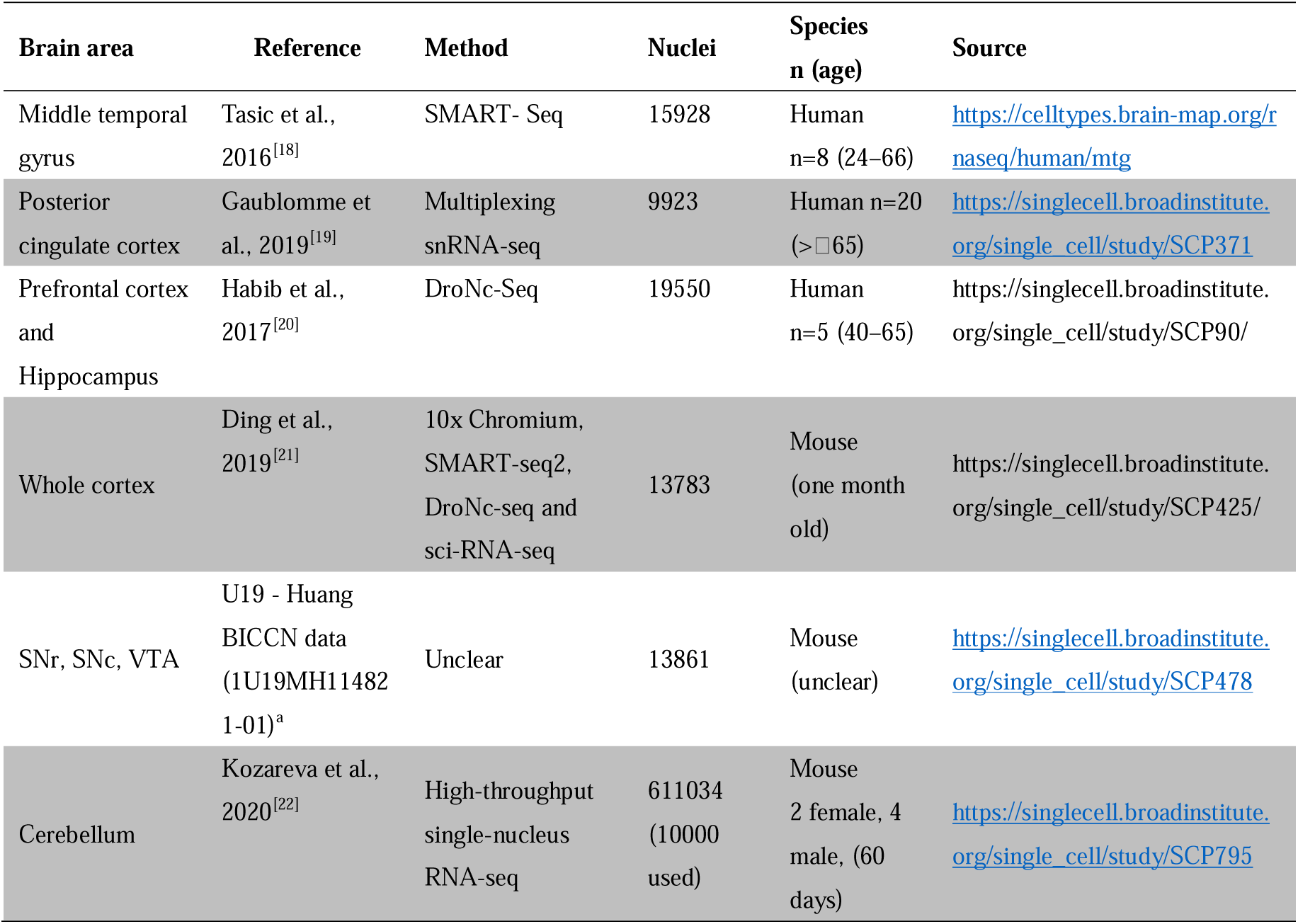
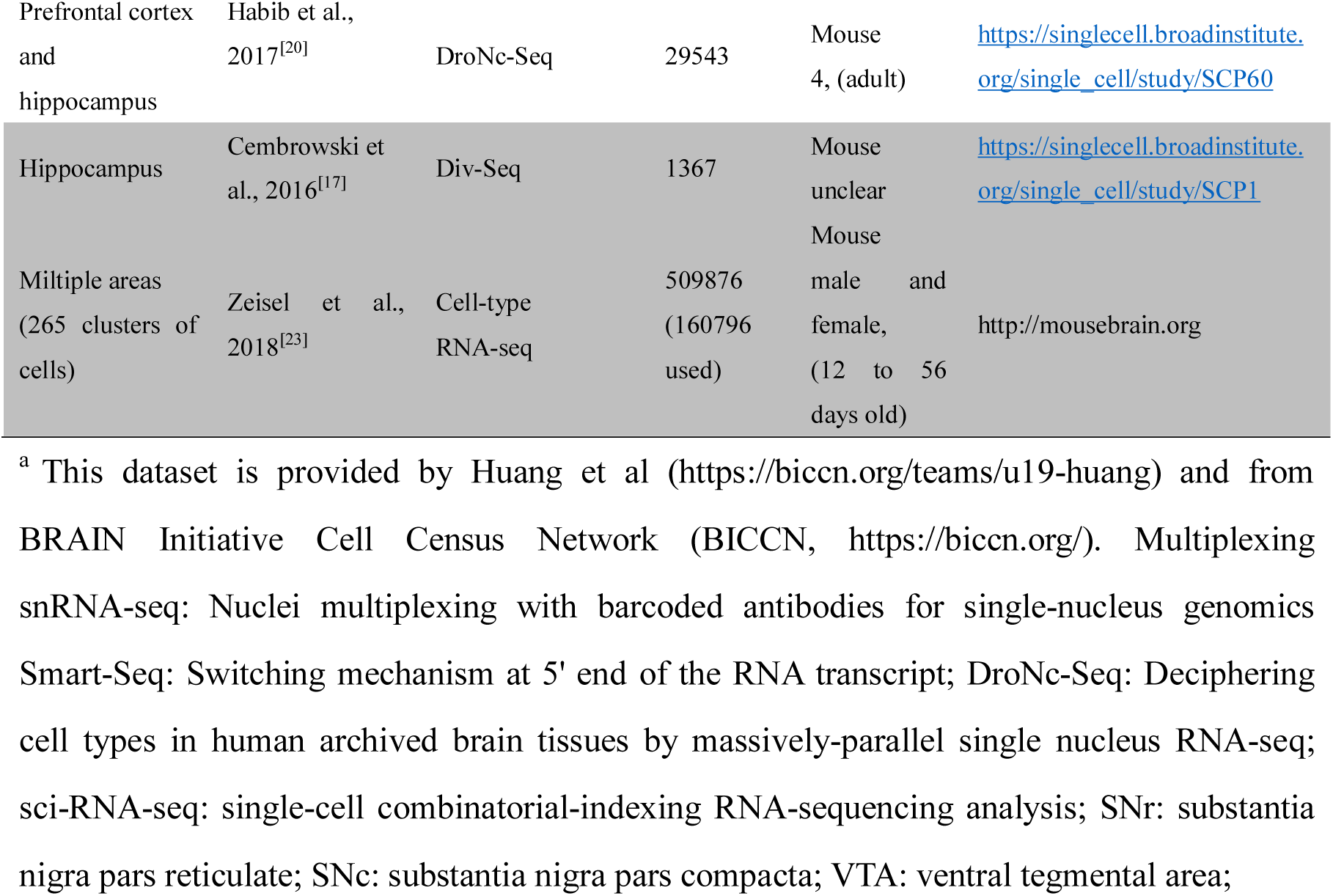
Summary of cell-type or single-cell sequencing databases for human and mouse brain.

### Spatial and cell-type distribution of Ace2 in mouse brain

Allen mouse brain atlas database (http://mouse.brain-map.org) was used to analyze the general spatial distribution of Ace2 in the mouse brain. Four cell-type sequencing datasets from hippocampus RNA-seq atlas, (https://hipposeq.janelia.org/), five single-cell sequencing datasets form Single Cell Portal database and one dataset Mouse brain atlas (http://mousebrain.org) were used for cell-type distribution of Ace2 in the mouse brain. The summary of the used five single-cell sequencing datasets was also shown in Table 2.

### Data processing and statistical analysis

Datasets were independently searched and analyzed by two authors (R.C. and J.Y.) and any disagreements were discussed and resolved by consensus with the corresponding author (Z.X.). The data from the Allen human brain atlas database and Allen Cell Types Database was exported to Microsoft Excel 2017 and GraphPad Prism 6.0 for further analysis. ACE2 expression data from the Allen human and mouse brain atlas database were also imported to Brain Explorer 2.0 software to get visualization. The data from the other databases were analyzed online.

To minimize false positive of expression as previous studies ^[24, 25]^, a gene with calculate counts per million (CPM), transcripts per million (TPM), unique molecular identifier (UMI) count or Fragments Per Kilobase Million (FPKM) > 1 were considered to be positive, which also is the same as log10 (CPM), log10 (TPM) or log10 (UMI) ≥ 0 as well as log10 (CPM+1), log10 (TPM+1), and log10 (UMI+1) ≥0.3. Besides, as Z score being greater than 2 corresponds to a p-value < 0.05, Z score of ACE2 expression > 2 were considered as high ACE2 expression in Allen human brain atlas data. Where applicable, data are expressed as median or mean. Interquartile range (IQR), 95%CI, range, and/or all sample points, were also provided if possible.

## Results

### The general expression of ACE2 in the human brain

According to the GETx portal database ^[15]^, compared with the lung, the general expression of ACE2 was lowe but not none in the brain (**Figure 1A**). The expression intensity of ACE2 was similar among different brain regions. Excepted for the lung (38 out of 578, 6.57%), the samples with log10(TMP+1) of ACE2 expression >0.3 were also found in amygdala (1 out of 152, 0.65%), anterior cingulate cortex (2 out of 176, 1.14%), caudate (3 out of 246, 1.22%), cortex (1 out of 255, 0.39%), frontal cortex (2 out of 209, 0.96%), hippocampus (4 out of 197, 2.03%), hypothalamus (3 out of 202, 1.49%), nucleus accumbens (1 out of 246, 0.41%), putamen (1 out of 205, 0.48%), spinal cord (cervical c-1, 4 out of 159, 2.52%), substantial nigra (5 out of 139, 3.60%), but none in cerebellum (0 out of 241) and cerebellar hemisphere (0 out of 215).

**Figure 1.**
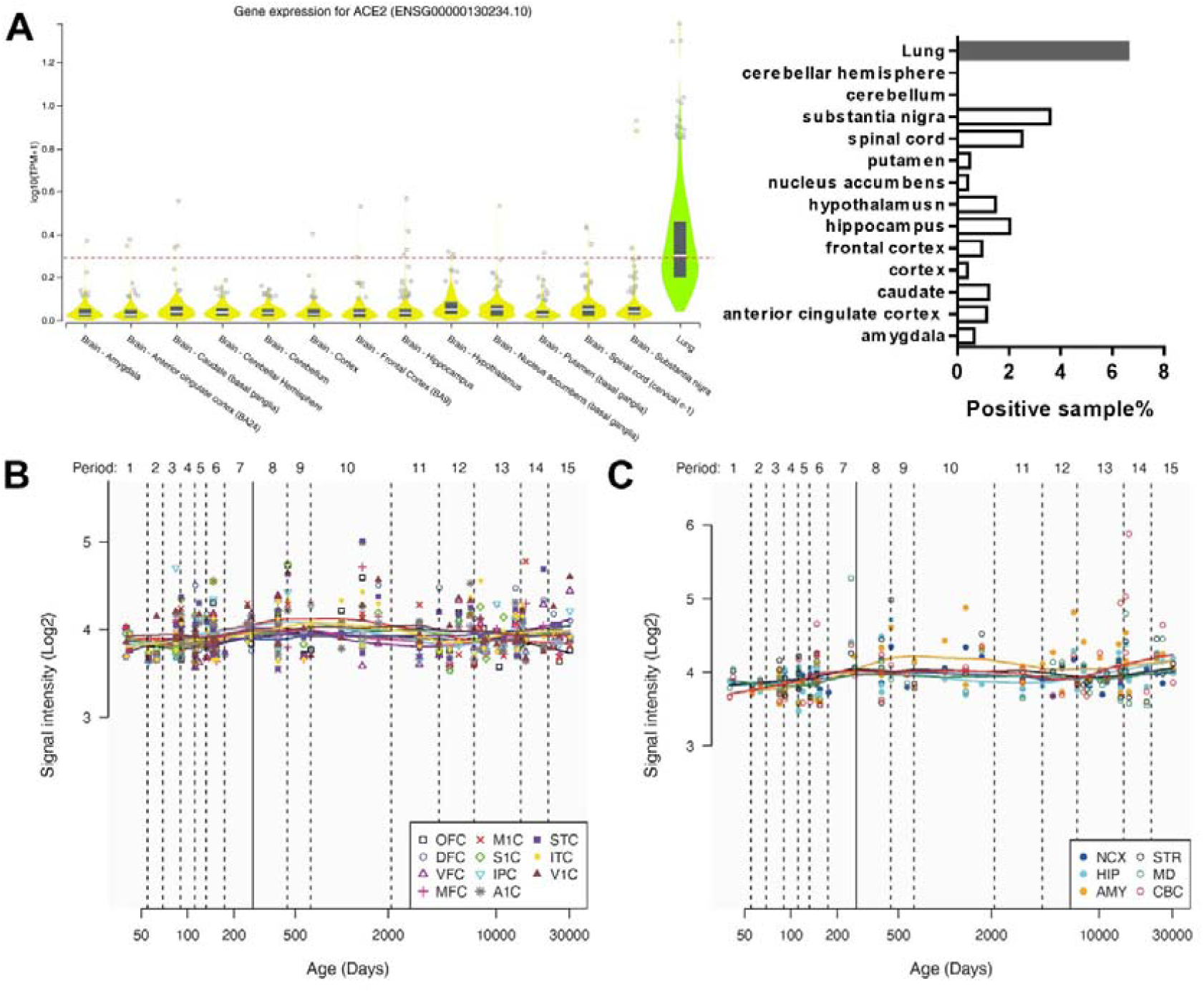
The general expression of ACE2 in different human brain areas. (**A**)The expression of ACE2 in different human brain areas and the lung according to GETx portal database ^[15]^. The dotted line means log10 (CPM+1) = 0.3, which means a threshold of the positive sample (>0.3 could be positive). (**B** and **C**) Change of intensity of ACE2 expression with age in different human brain areas according to the Human Brain Transcriptome database ^[13]^. Data are expressed as the median, interquartile range (IQR), and all sample points in **A**, while data are expressed as mean and all sample points in **B** and **C**.

Besides, according to the Human Brain Transcriptome database ^[13]^, the expression of ACE2 was also similar among cortex and other brain regions, which may not change a lot with the age (**Figure 1B and C**).

### Spatial distribution of ACE2 in the human brain

By providing slice images, Allen Human Brain Atlas may prove a more detailed Spatial distribution of ACE2 in the human brain than the GETx portal database and Human Brain Transcriptome database ^[14]^. Two microarray datasets using the different probes of ACE2 (A_23_P252981, CUST_16267_PI416261804) were found and used in Allen Human Brain Atlas (Http://human.brain-map.org/microarray/search). By analyzing the intensity of ACE2 expression, we found six brain areas with a maximal z score of ACE2 expression >2.0 in both probe datasets, 15 brain areas with a maximal z score of ACE2 expression >2.0 only in probe1 datasets, and 9 which represents ACE2 expression in probe1 datasets. Z score of ACE2 expression >2.0 means that these brain areas are 2 standard deviations greater than the mean, which corresponds to a p-value < 0.05 (**Figure 2A and B**).

**Figure 2.**
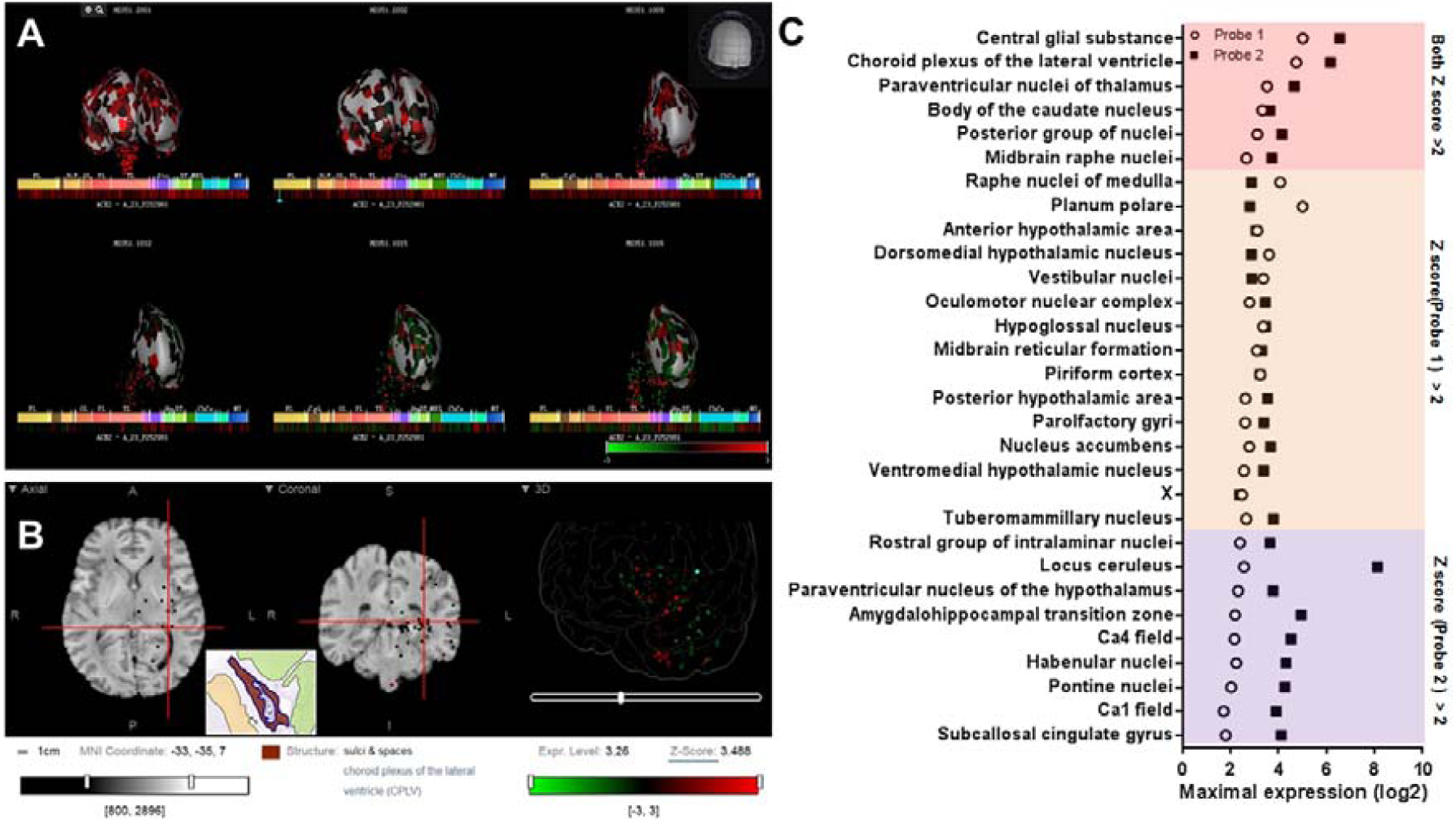
Spatial distribution of ACE2 in the human brain according to Allen Human Brain Atlas. (**A**) 3D view of the expression of ACE2 in different human brain areas based on the data of probe A_23_P252981 (probe 1). (**B**) Planar view of the expression of ACE2 in the choroid plexus of the lateral ventricle. The inset shows a sampled area of the choroid plexus of the lateral ventricle (the dark area in the inset). (**C**) log2 intensity of ACE2 expression in the 30 brain areas with Z score > 2.0 in at least one probe dataset (probe 1: A_23_P252981; probe 2: CUST_16267_PI416261804). All data were generated from the Allen Human Brain Atlas (Http://human.brain-map.org/microarray/search). Images in A and B directly from the Allen Human Brain Atlas (© 2010 Allen Institute for Brain Science. Allen Human Brain Atlas. Available from: human.brain-map.org).

By pooled the expression value of the two datasets, we further found 23 brain areas with z score >1.0 and four of them with z score >2.0 (**Figure 3A**). The distribution view of ACE2 expression in the human brain according to the pooled expression data was showed in **Figure 3B**.

**Figure 3.**
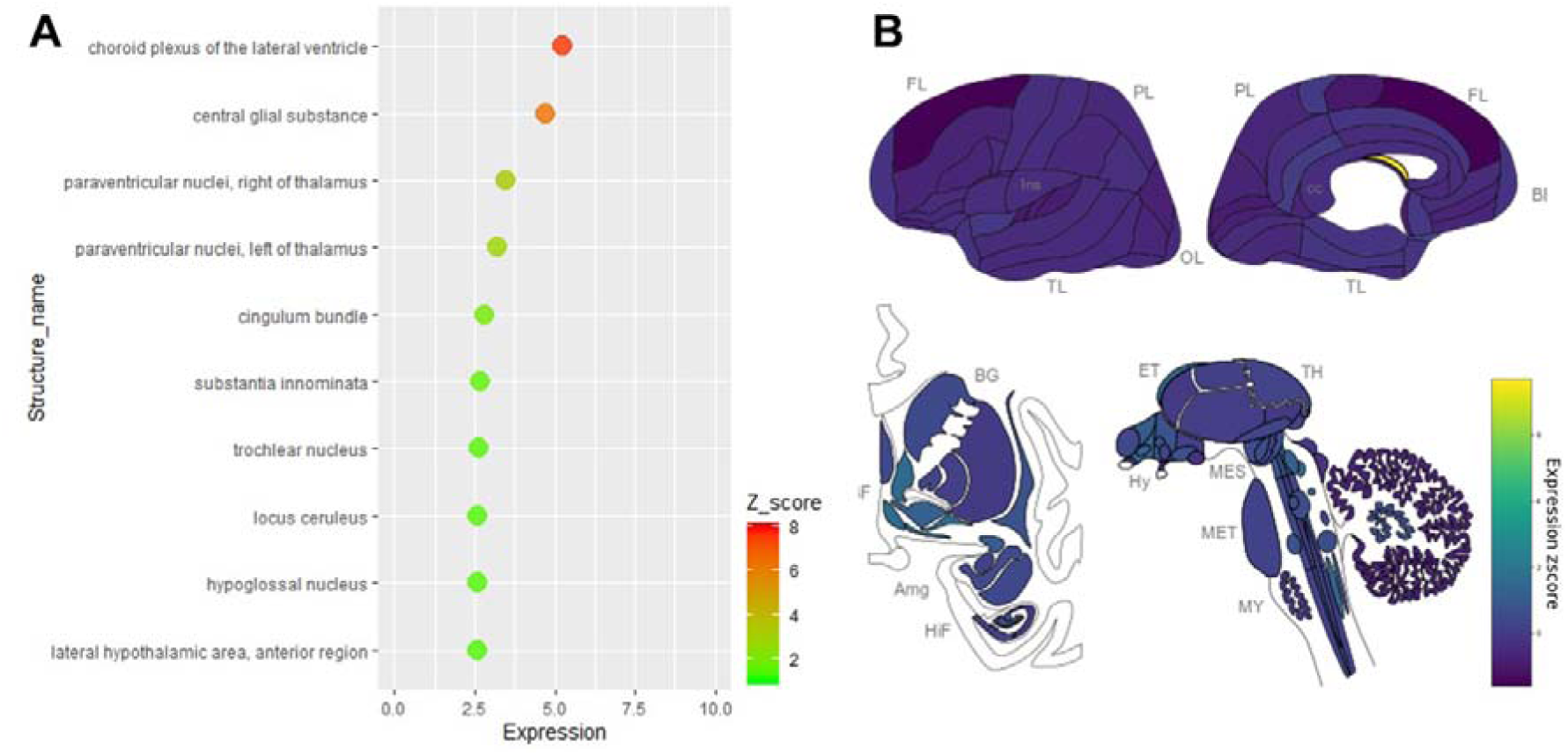
Spatial distribution of ACE2 in the human brain according to the pooled data of Allen human Brain Atlas.. (**A**) the expression intensity and Z-score of ACE2 in the top 10 brain areas of pooled expression data of the two probes. (**B**) the distribution view of ACE2 expression in the human brain according to the pooled expression data of the two probes. Original data are available from Allen Human Brain Atlas (http://human.brain-map.org/microarray/search). Brain region abbreviations: frontal lobe (FL); parietal lobe (PL); temporal lobe (TL); occipital lobe (OL); basal forebrain (BF); basal ganglia (BG); amygdala (AmG); hippocampal formation (HiF); epithalamus (EP); thalamus (TH); hypothalamus (Hy); mesencephalon (MES); metencephalon (MET) and myelencephalon (MY).

### Cell-type distribution of ACE2 in the human brain

We further collected and analyzed single-cell sequencing data, which may provide all mRNAs present in every single cell of tested brain tissue. By analyzing the single-cell sequencing data of human middle temporal gyrus (https://celltypes.brain-map.org/rnaseq/human/mtg^[18]^), human posterior cingulate cortex (https://singlecell.broadinstitute.org/single_cell/study/SCP371), and archived human prefrontal cortex and hippocampus samples (https://singlecell.broadinstitute.org/single_cell/study/SCP90/), the expression of ACE2 is relatively high in human middle temporal gyrus (totally 2.00% ACE2 positive cells, 309 out of 15603) and posterior cingulate cortex (totally 1.38% ACE2 positive cells, 133 out of 9635) samples, but it is very low in the archived human prefrontal cortex (no ACE2 positive cells) and hippocampus samples (totally 0.21% ACE2 positive cells, 2 out of 9530).

For the cell types, most of the ACE2 positive cells are neurons in both human middle temporal gyrus and posterior cingulate cortex, especially excitatory neurons (72.1% for middle temporal gyrus and 66.1% for posterior cingulate cortex, **Figure 4A and B**) and interneurons (22.4% for middle temporal gyrus and 9.8% for posterior cingulate cortex, **Figure 4A and B**). The percentage of positive cells in excitatory neurons was also the highest among cell types (2.14% for middle temporal gyrus and 2.18% for the posterior cingulate cortex, **Figure 4C**). The details of the cell-type distribution of ACE2 in the human brain were shown in **Supplementary Figures 1 and 2**.

**Figure 4.**
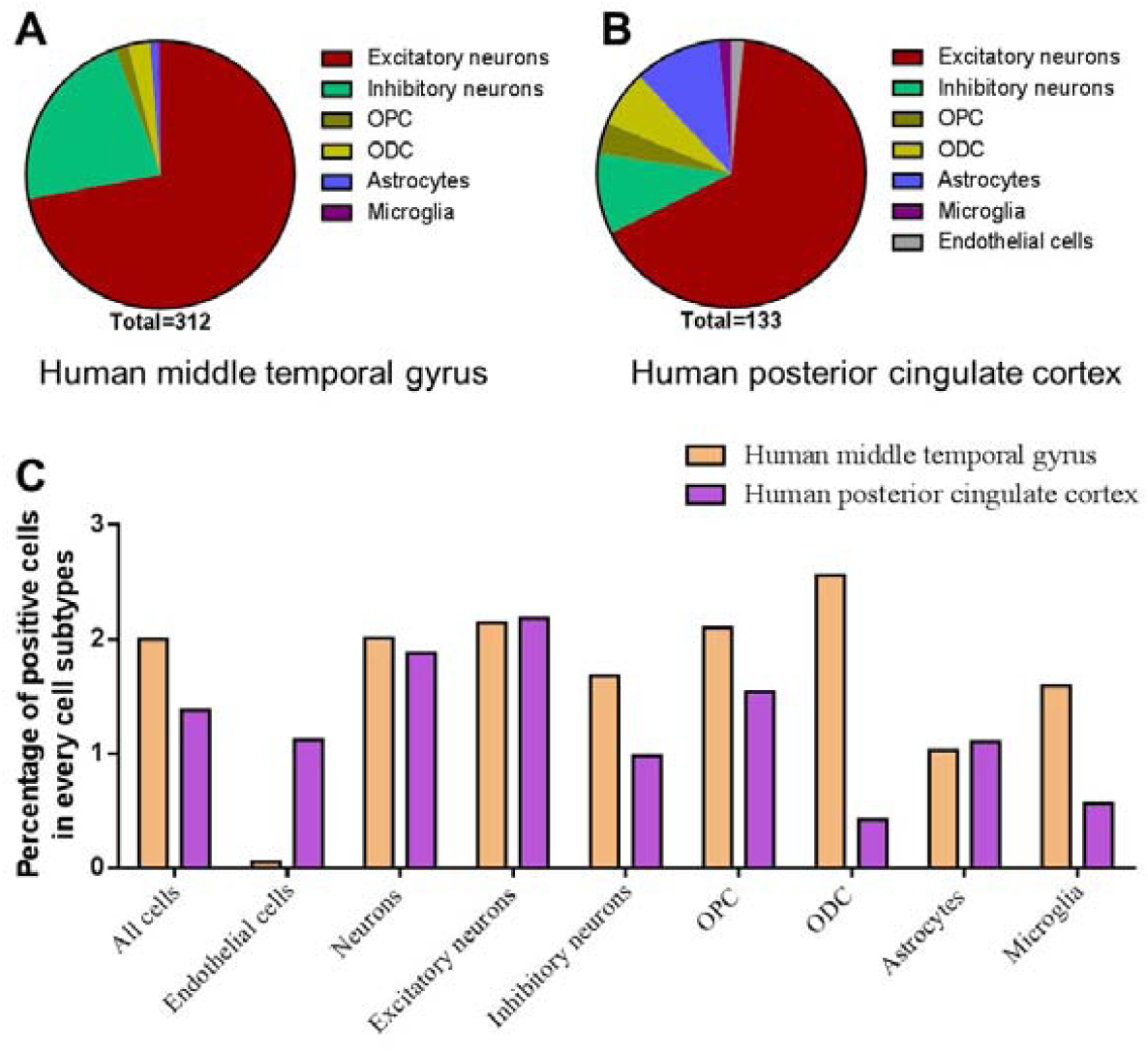
Cell-type distribution of ACE2 in the human brain. **(A)** The Cell-type proportion of total positive cells in the human middle temporal gyrus. (B) The Cell-type proportion of total positive cells in the human posterior cingulate cortex. **(C)** The percentage of positive cells in different cell subtypes in the human middle temporal gyrus and posterior cingulate cortex. Original data are from https://celltypes.brain-map.org/rnaseq/human/mtg and https://singlecell.broadinstitute.org.

### Spatial and Cell-type distribution of Ace2 in the mouse brain

As shown in **Figure 5**, we additionally analyzed the spatial distribution of Ace2 in the mouse brain based on the Allen Mouse Brain Atlas (http://mouse.brain-map.org/gene/show/45849; Lein et al., 2007). Similar to the human brain, we found the Ace2 expression is relatively high in the choroid plexus of lateral ventricle, substantia nigra pars reticulata (SNr), and some cortical areas (such as the piriform cortex). We additionally found the ACE2 expression is also relatively high in the olfactory bulb, while it was very low in most hippocampal areas.

**Figure 5.**
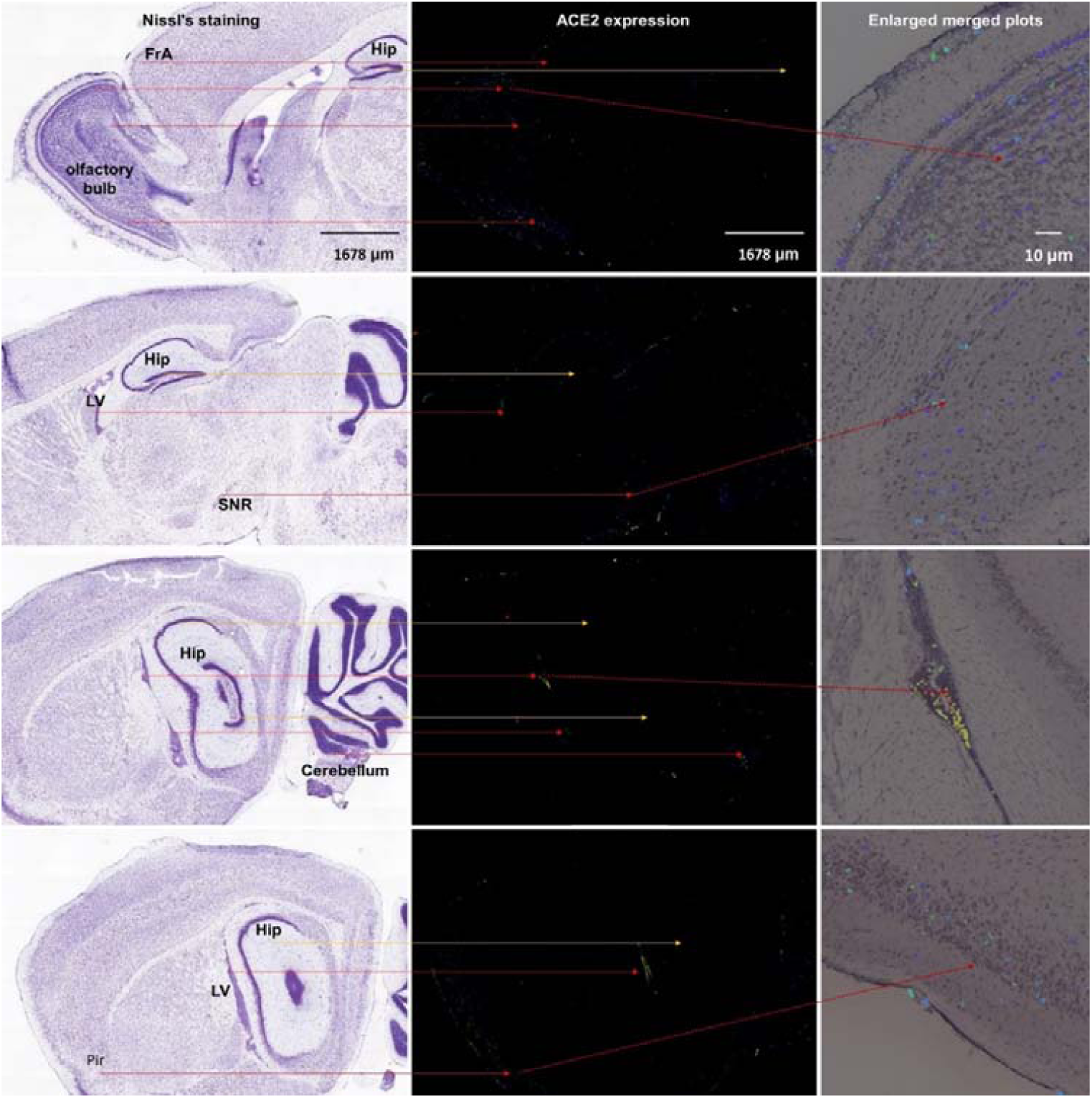
Spatial distribution of ACE2 in mouse the brain. The left column means Nissl’s staining of mouse brain slice, the middle column means the ACE2 expression by antisense, the right column means the enlarged merged plots form Brain Explorer 2.0 software. Hip: hippocampus; LV: lateral ventricle; SNR: substantia nigra pars reticulate. Pir: piriform cortex. All images are from the Allen Mouse Brain Atlas (© 2004 Allen Institute for Brain Science. Allen Mouse Brain Atlas. Available from https://mouse.brain-map.org/).

We further analyze the single-cell sequencing data of multiple mouse cortex (https://singlecell.broadinstitute.org/single_cell/study/SCP425/), archived mouse brain samples (https://singlecell.broadinstitute.org/single_cell/study/SCP60), mouse hippocampus (https://singlecell.broadinstitute.org/single_cell/study/SCP1;), substantia nigra pars reticulate (SNr), substantia nigra pars compacta (SNc) and ventral tegmental area (VTA) areas (https://singlecell.broadinstitute.org/single_cell/study/SCP478), and cerebellum (https://singlecell.broadinstitute.org/single_cell/study/SCP795). Similar to human data, the expression of Ace2 is relatively high in multiple mouse cortex samples (totally 0.84% Ace2 positive cells, 84 out of 10000), but it is relatively low in mixed mouse prefrontal cortex and hippocampus samples (totally 0.28% Ace2 positive cells, 38 out of 13313) and mouse hippocampus samples (totally 0.1% Ace2 positive cells, 2 out of 1188). Besides, the expression of Ace2 is also found in 0.68% cell in mixed SNr, SNc, and VTA areas as well as 0.52% in the cerebellum.

For the cell-types distribution, most of the Ace2 positive cells are endothelial cells in the multiple mouse cortex samples, the mixed prefrontal cortex and hippocampus samples, the mixed SNr, SNc and VTA areas and the cerebellum (**Figure 6**). The percentage of positive cells in endothelial cells was the highest among different cell types in most mouse brain. Of note, the neurons also account for a relatively large proportion of total Ace2 positive cells in multiple cortex samples (48.8%) and mixed prefrontal cortex and hippocampus samples (15%), the mixed SNr, SNc, and VTA areas (10.5%) and the cerebellum (7.69%) in mouse brain. The details of the cell-type distribution of Ace2 in the mouse brain are shown in **Supplementary Figure 3**. Similarly, in a comprehensive atlas of the mouse nervous system, of the 265 cell clusters, four had an expression Z score > 2 (three pericyte clusters and a cluster of arterial endothelial cells[23]).

**Figure 6.**
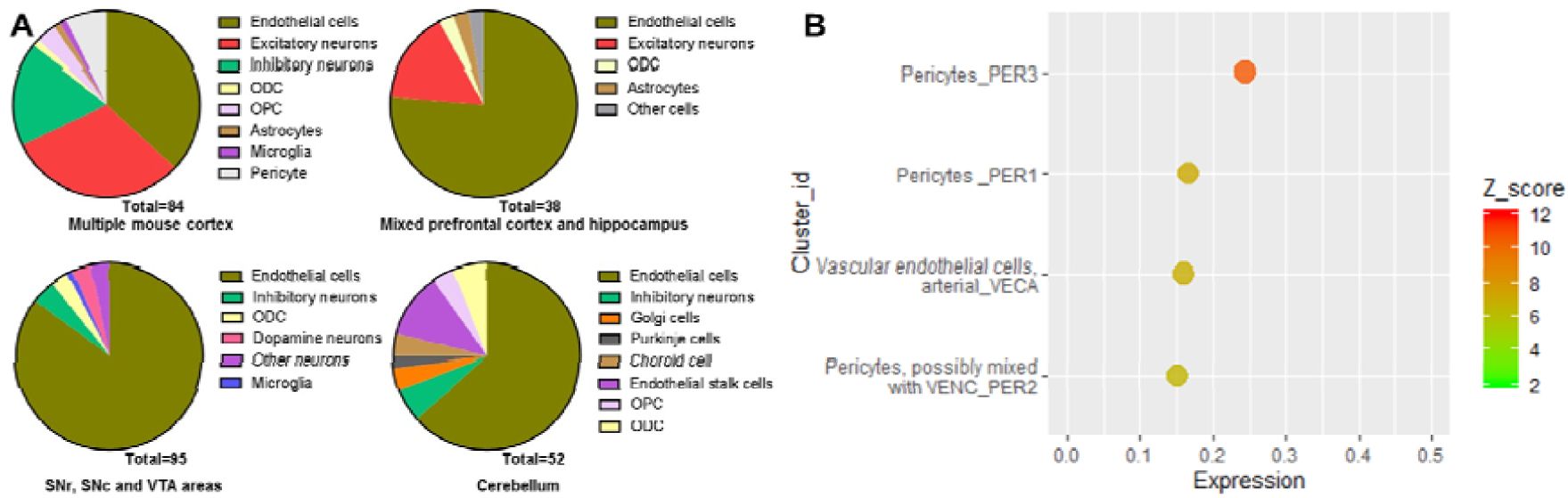
Cell-type distribution of Ace2 in mice brain. (A) Cell-type distribution of Ace2 according to Single Cell Portal database (https://singlecell.broadinstitute.org/) ; (B) Top cell-type cluster expression of Ace2 according to Mouse brain atlas database (http://mousebrain.org).

In addition, as the inconsistent results were found in the hippocampus based on the single-cell sequencing, we additionally analyzed the expression of Ace2 in a cell-type sequencing database of the mouse hippocampus, the Hippocampus RNA-seq atlas database (https://hipposeq.janelia.org; Cembrowski et al., 2016). As shown in **Supplementary Figure 4**, (1) Ace2 expression was only found in intermediate and ventral areas of the hippocampus with the mean FPKM<1; (2) Ace2 expression were found in ventral pyramidal cells but not in dorsal CA1 pyramidal cells, dorsal CA3 pyramidal cells, dorsal CA2 pyramidal cells, dorsal DG granule cells, dorsal DG mossy cells, ventral CA3 pyramidal cells, and ventral DG granule cells; (3) Ace2 expression were not found in PV-positive or SST-positive interneurons in the hippocampus; (4) Ace2 expression were not found in any neurons that project to post-subiculum, NAc or amygdala.

## Discussion

ACE2 is an important entry receptor for SARS-CoV-2 and SARS-CoV infecting host organs ^[6]^. Though the infection of SARS-COV in the brain was reported in the past ^[26, 27]^, the distribution of ACE2 in the brain is still unclear. Here we mainly found: (1) The expression of ACE2 was relatively high in several specific brain areas in human; (2) the expression of ACE2 mainly located in many neurons (both excitatory and inhibitory neurons) in human middle temporal gyrus and posterior cingulate cortex, but the ACE2-expressing cells were none in PFC and very few in the hippocampus; (3) Except for the additional expression of ACE2 in the olfactory bulb areas for spatial distribution and the pericytes and endothelial cells for cell-type distribution analysis, the main distribution figure of ACE2 in mouse brain was similar to that in human. Thus, our results reveal an outline of ACE2 distribution in the human and mouse brain, which supports the hypothesis that the SARS-CoV-2 is capable to infect the brain and may result in CNS symptoms in COVID-19 patients ^[28]^.

SARS-CoV-2 shares a 77.2% amino acid identity, 72.8% sequence identity, and high structural similarity to previous SARS-CoV ^[29, 30]^. Similar to SARS-CoV, experimental affinity measurements show a high affinity of the receptor-binding domain of SARS-CoV-2 and ACE2 ^[30, 31]^. Here we found the percent of ACE2 positive samples were found in most brain areas, especially substantial nigra was almost comparable to that of the lung (3.60% vs 6.57%), though the total expression in the brain seems much lower than that in the lung. According to Allen brain atlas, the total expression of ACE2 seems not to change with age. Six brain areas with a Z score of ACE2 expression >2 in both probe data and four brain areas were Z score of ACE2 expression >2 in pooled data. Some brain nuclei are very important for the normal brain functions, including the paraventricular nuclei of the thalamus (involved in the control of wakefulness, feeding, appetitive motivation, drug addiction, regulation of stress and negative emotional behavior, and epilepsy^[32, 33]^), the raphe nuclei (the main serotoninergic nuclei in the brain^[34]^), and tuberomammillary nucleus (the main histaminergic nuclei in the brain ^[35]^). Thus, our results highlight the importance of spatial distribution rather than the general total expression of ACE2 in the brain. These results may provide some clues to further study on the brain infection of SARS-CoV-2 in the COVID-19 patients, and suggesting SARS-CoV-2 might be able to result in serious CNS symptoms in COVID-19 patients (if it could infect these important brain areas by binding ACE2). Brain imaging and long-term follow up may be needed in COVID-19 patients for the possibility of SARS-CoV-2 brain infection and the following brain disorders.

The routes or pathways for SARS-CoV and novel SARS-CoV-2 entering the brain are still unclear. According to experiments in mice transgenic for human ACE2, intranasally given SARS-CoV may enter the brain by olfactory nerves ^[36]^. Consistent with this, we found the expression of ACE2 in the olfactory bulb is higher than that in most other cortexes (Figure 3). In the human brain, we found the piriform cortex, a brain area directly connected with olfactory bulb, was ACE2 high-expression. Though no ACE2 expression data of olfactory bulb in humans was available, our results indirectly support the hypothesis that SARS-CoV-2 might enter the human brain by olfactory nerves. On the other hand, we additionally found the central glial substance and choroid plexus of the lateral ventricle were with very high ACE2 expression (Z score > 5) in the human brain. Relatively high expression of ACE2 in the choroid plexus of the lateral ventricle was also found in the mouse brain in the current study. The central glial substance refers to an area of grey matter surrounding the central canal, which carries CSF and helps to transport nutrients to the spinal cord ^[37]^; Besides, the choroid plexus of ventricles is an important brain area for the generation of CSF [38], the main location of the blood-cerebrospinal fluid barrier ^[39]^, and a crucial gateway for immune cells entering the brain ^[40]^. Recently, the SARS-CoV-2 has also been found in cerebrospinal fluid (CSF) samples from a 56-year-old COVID-19 patient by genetic sequencing in China (http://www.ecns.cn/news/society/2020-03-05/detail-ifzuhesu4119860.shtml). SARS-CoV-2 may also infect the brain in a 24-year-old male patient ^[25]^. Thus, our results suggest the high expression of ACE2 in the central glial substance and ventricles may provide another potential pathway for the SARS-CoV-2 or SARS-CoV entering CSF and/or spreading around the brain.

Single-nucleus RNA-seq provides a high resolution of cellular gene-expression of each cell ^[41]^. According to single-nucleus RNA-seq data, we further found that ACE2 located in many neurons (especially excitatory neurons) and some non-neuron cells (especially astrocytes and ODCs) in both posterior cingulate cortex and middle temporal gyrus, where the expression level of ACE2 seems low under normal conditions. Here, the highest number of ACE2 positive cells were excitatory neurons, which may be projection neurons that make up many important brain networks. For example, excitatory neurons in the posterior cingulate cortex may project dense connections to the hippocampal formation and parahippocampal cortex, which are related to emotion and memory ^[42]^. On the other hand, though the positive cell number of inhibitory neurons is lower than excitatory neurons, the percentage of positive cells in some sub-types inhibitory neuron were comparable or slightly low compared with excitatory neurons. Inhibitory neurons are also crucial for normal brain function ^[43]^. For example, the neurons in SNr are mainly inhibitory GABAergic neurons, which is one important note in the neural circuits that contribute to epilepsy ^[33]^. Besides, we also found some dopaminergic neurons and cerebellar cells in the mouse brain are also ACE2-positive. Thus, our results may help to explain the previous finding that SARS-CoV particles are mainly located in the neurons in the brain samples from SARS patients ^[26]^, and suggest SARS-CoV-2 may also invade many neurons in the human brain and hence contribute to the CNS symptoms in COVID-19 patients.

In addition, single-nucleus RNA-seq data showed that the ACE2 was found in very few endothelial cells and pericytes in the human brain, while endothelial cells and pericytes were the main Ace2-expression cell types in the mouse brain. Of note, previous microarray data shows ACE2 expressed in the human endothelium ^[9]^. Endothelial cells are the main component of blood vessel endothelium. Thus, one possible reason for these differences could be that most blood vessels may be excluded from the tested human brain tissues while they may be hard to be removed in the mouse brain. Both endothelial cells and pericytes are closely related to the blood-brain barrier integrity ^[44]^. Endothelial cells and pericytes could also build blocks of the neurovascular unit together with astrocytes and neurons ^[45]^. Thus, endothelial cells and pericytes may help the SARS-CoV-2 to enter the brain by crossing the BBB and also infect the neurovascular unit, which may contribute to the cerebrovascular events in COVID-19 patients ^[46]^. Taken together, as there is no enough evidence supporting the ACE2 high expression in endothelial cells and pericytes in the human brain, further studies remain needed. The CSF biomarkers of pericytes injury and blood-brain barrier integrity, such as sPDGFRβ and albumin respectively ^[47]^, may be needed to detect whether SARS-CoV-2 damaged the BBB in COVID-19 patients in future studies. Besides, such a potential difference between humans and mice should be noted when using mouse models for the SARS-CoV-2 study.

## Conclusions

Our results reveal an outline of ACE2 or Ace2 distribution in the human and mouse brain, which indicates the brain infection of SARS-CoV-2 might be capable to infect the brain and result in CNS symptoms in COVID-19 patients. The finding of high ACE2 expression in central glial substance and brain ventricles suggests two potential novel routes for the SARS-CoV-2 entering the CSF and/or spreading around the brain. Besides, the differences of ACE2/Ace2 distribution between humans and mice may be also useful to further “bench to bedside” translational studies regarding SARS-CoV-2. Our results may help to bring light to the brain infection of the present novel SARS-CoV-2 and previous SARS-CoV. Further studies are warranted to confirm our results and related predictions.

## Acknowledgments

We would like to thank the Allen Institute for Brain Science (https://alleninstitute.org/), BRAIN Initiative Cell Census Network (BICCN, https://biccn.org/), GTExportal database (https://www.gtexportal.org), Human Brain Transcriptome database (https://hbatlas.org), Single Cell Portal database (https://singlecell.broadinstitute.org), and Mouse brain atlas database (http://mousebrain.org).

## Funding

This work was funded by the Natural Science Foundation of Zhejiang Province (LEZ20H190001) and the National Natural Science Foundation of China (81673623). Partly supported by and the Foundation of Zhejiang Chinese Medical University (Q2019Y02, ZYX2018002).

## Compliance with ethical standards

### Ethics approval

Not Applicable. All the data in this study are from a publicly available online database.

### Competing interests

None declared.

### Authors’ contributions

Z.X. and C.W. designed the study. R.C., K.W., and J.Y. performed the search and analysis. D.H. and L.F. contribute to the data analysis for Figure 1 and 2. Z.X. and K.W. checked the analyzed data. Z.X. wrote the manuscript in consultation with J.Y., Z.C., C.W., D.H., and L.F..

## Notes

### Competing Interest Statement

The authors have declared no competing interest.

### Summary of Updates

With the help of Derek Howard and Leon French, we further analyzed a new database (Mouse brain atlas database (http://mousebrain.org). We revised Figure 6. All Supplementary Figures were moved to the supplementary file. Thank you very much.

