## Supplementary Figure for "The spatial and cell-type distribution of SARS-CoV-2 receptor ACE2 in human and mouse brain"

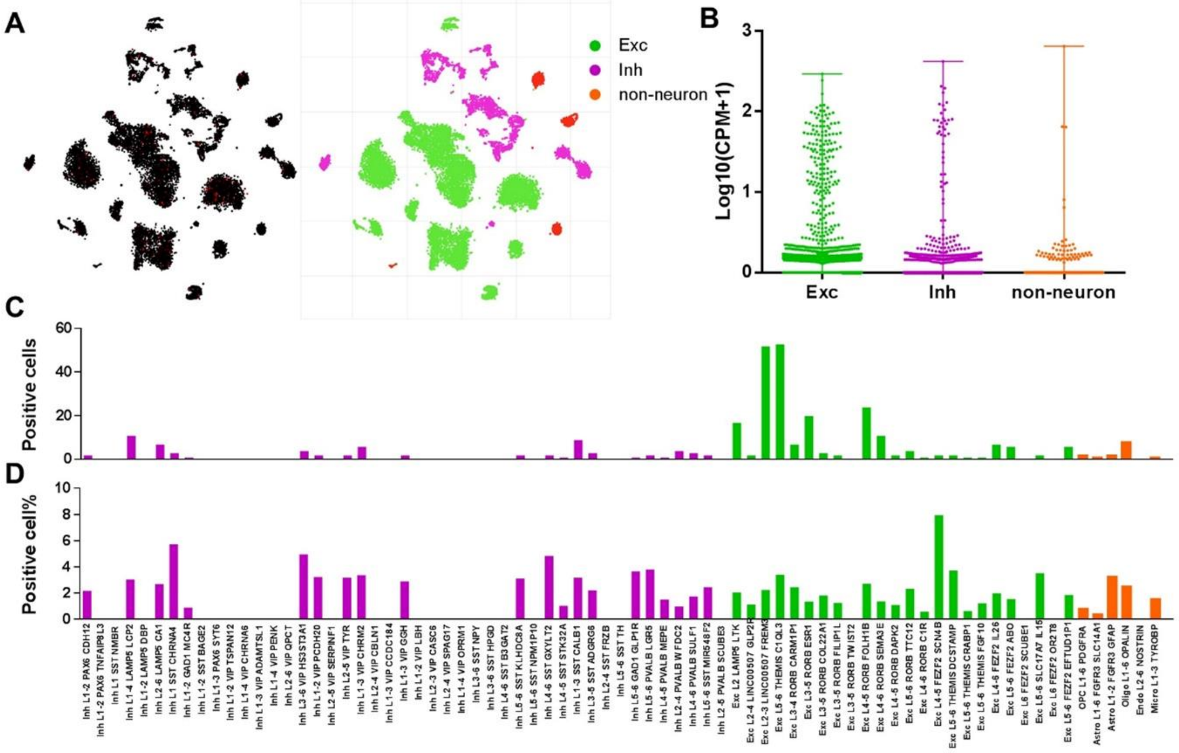


**Supplementary Figure 1. ACE2 expression in the human middle temporal gyrus (related to Figure 4).** (**A**) The expression of ACE2 in the cells of the human middle temporal gyrus. Left: Red means potential ACE2-positive cells, while black means ACE2-negative cells; Right: the annotation map of cell clusters. (**B**) The expression of ACE2 in brain cell subtypes. (**C**) The number of ACE2-positive cells in each sub-cluster of brain cells. (**D**) The percentage of ACE2-positive cells in each sub-cluster of brain cells. Cells with log10 (CPM+1) >0.3 were considered as positive cells. CPM: calculate counts per million; Inn: Inhibitory neuron; Exc: Excitatory neuron; Oligo: oligodendrocyte; Astro: astrocytes; Micro: microglia. Original data are from https://celltypes.brain-map.org/rnaseq/human/mtg. Images in A were directly generated by the RNA-Seq Data Navigator from Allen Cell Types Database (© 2015 Allen Institute for Brain Science. Allen Cell Types Database. Available from https://celltypes.brain-map.org/). Data are expressed as median, range, and all sample points in B.

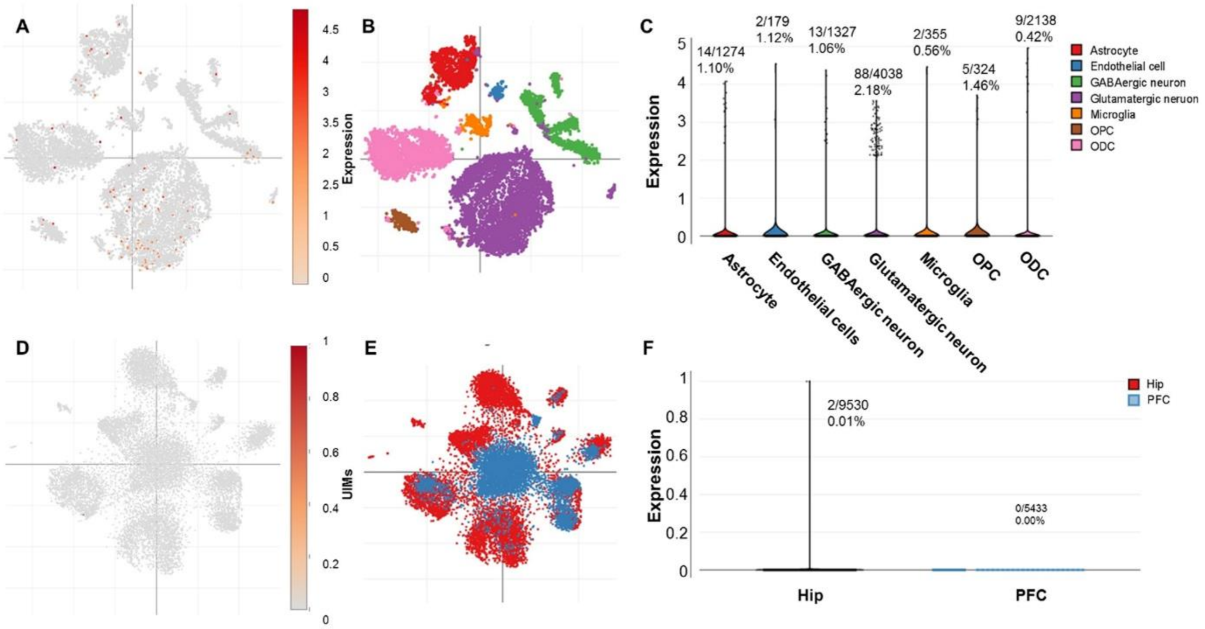


**Supplementary Figure 2. Expression of ACE2 in the human posterior cingulate cortex, prefrontal cortex, and hippocampus (related to Figure 4).** (**A**) Expression of ACE2 in the human posterior cingulate cortex. (**B**) The annotation map of cell clusters. (**C**) Expression of ACE2 in brain cell subtypes. (**D**) Expression of ACE2 in human prefrontal cortex and hippocampus. (**E**) The annotation map of brain regains, red points mean cells from the prefrontal cortex, and blue points mean cells from the hippocampus. (**F**) Expression of ACE2 in human prefrontal cortex and hippocampus. Original data are available from https://singlecell.broadinstitute.org. Data are expressed as median, range, and all sample points in C and F.

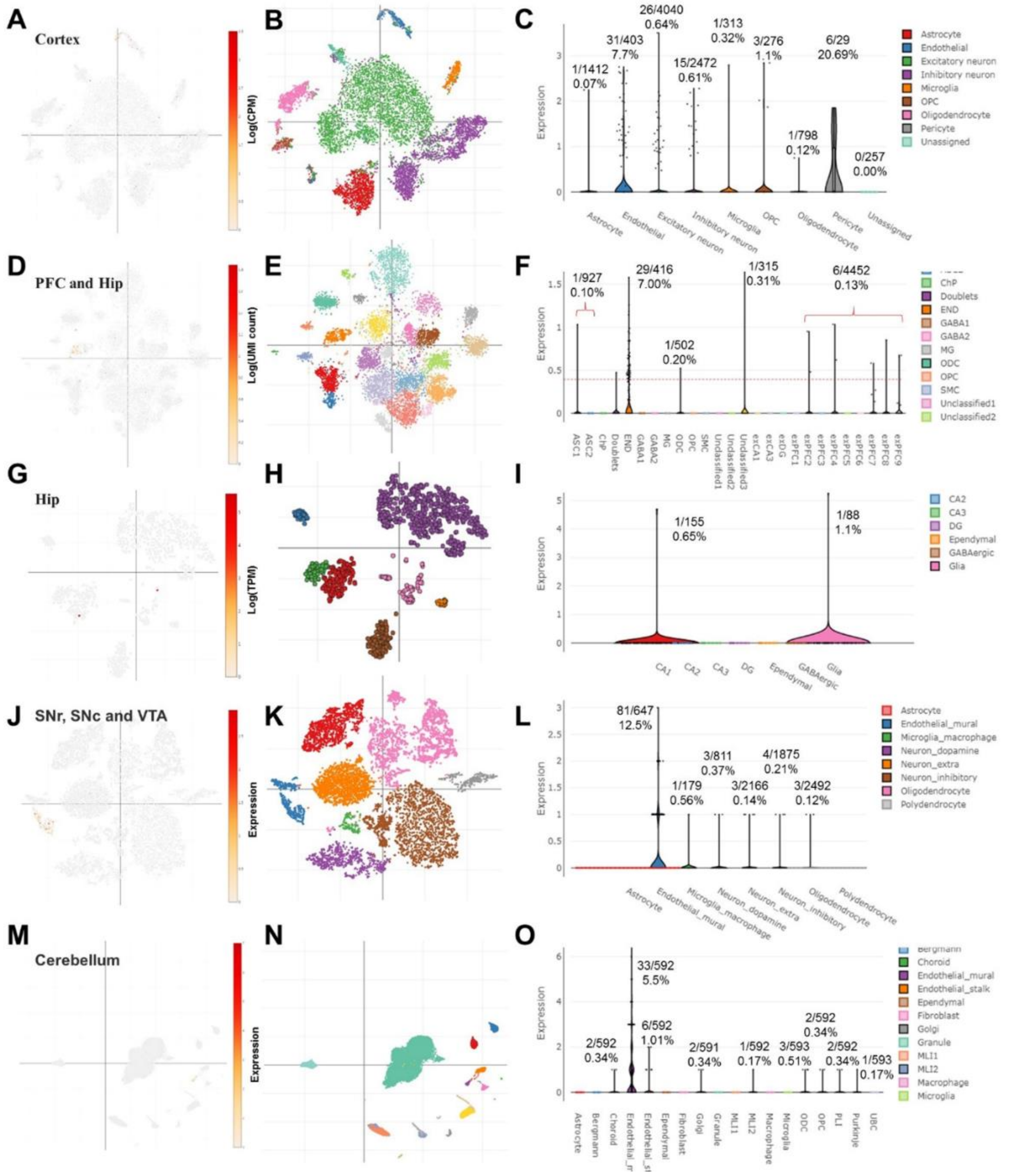


**Supplementary Figure 3. Cell-type distribution of Ace2 in the mouse brain (related to figure 6).** (**A-C**) Cell-type distribution of Ace2 in the mouse cortex. (**A**): Ace2 expression map; (**B**): the annotation map; (**C**) distribution plot. (**D-F**) Cell-type distribution of Ace2 in the mouse prefrontal cortex and hippocampus. (**D**): Ace2 expression map; (**E**): the annotation map; (**F**) distribution plot. (**G-I**) Cell-type distribution of Ace2 in mouse hippocampus. (**G**): Ace2 expression map; (**H**): the annotation map; (**I**) distribution plot. (**J-L**) Cell-type distribution of Ace2 in mouse SNr, SNc, and VTA. (**J**): expression map; (**K**): the annotation map; (**L**) distribution plot. (**M-O**) Cell-type distribution of Ace2 in mouse cerebellum. (**M**): Ace2 expression map; (**N**): the annotation map; (**O**) distribution plot. ACS: astrocyte; ChP: choroid plexus cell; END: endothelial cells; Hip: hippocampus; MG: microglia; MLI: molecular layer interneuron; ODC: oligodendrocyte; OPC: Oligodendrocyte progenitor cell. PFC: prefrontal cortex; SMC: smooth muscle cell; SNr: substantia nigra pars reticulate; SNc: substantia nigra pars compacta; VTA: ventral tegmental area; UBC: unipolar brush cells. Data are expressed as median, range and all sample points in **C, F, I, L, and O**. Original data are available from https://singlecell.broadinstitute.org.

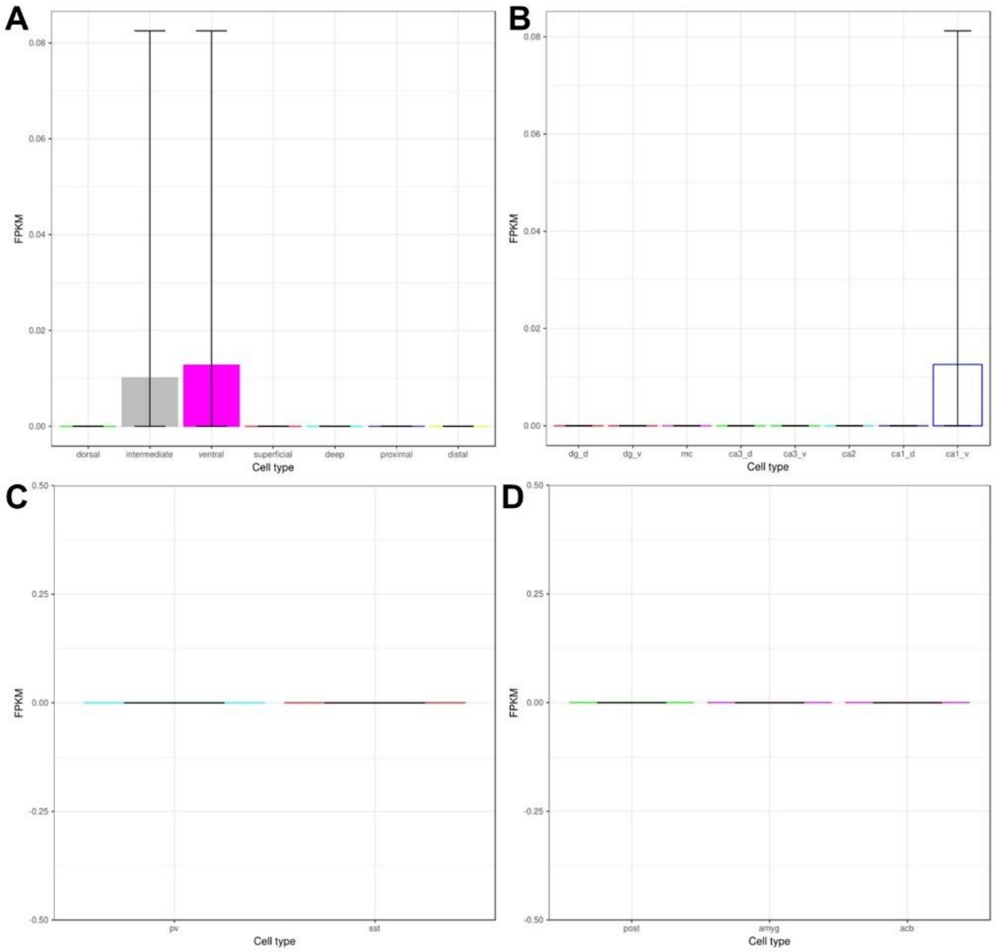


**Supplementary Figure 4. Spatial and cell-type distribution of ACE2 in the mouse hippocampus (related to figure 6).** (**A**) Spatial distribution of ACE2 in mouse hippocampus. (**B**) Spatial distribution of ACE2 in mouse hippocampal pyramidal cells and granule cells. dg_d: Dorsal DG granule cell; dg_v Ventral DG granule cell; mc: Dorsal DG mossy cell; ca3_d: Dorsal CA3 pyramidal cell; ca3_v: Ventral CA3 pyramidal cell; ca2: Dorsal CA2 pyramidal cell; ca1_d: Dorsal CA1 pyramidal cell; ca1_v: Ventral CA1 pyramidal cell (**C**) Cell-type distribution of ACE2 in mouse hippocampal interneurons. Pv: parvalbumin-expressing neurons; sst: somatostatin-expressing neurons. (**D**) Distribution of ACE2 in mouse hippocampal neurons that project to post-subiculum, nucleus accumbens or amygdala. post: post-subiculum projecting neuron; amyg: amygdala projecting neuron. NAc; nucleus accumbens projecting neuron. FPKM: Fragments Per Kilobase Million. Data are expressed as mean and 95%CI. Original data are available from: https://hipposeq.janelia.org (Cembrowski et al., 2016)
